# Quantification of lactoyl-CoA by liquid chromatography mass spectrometry in mammalian cells and tissues

**DOI:** 10.1101/2020.03.15.992859

**Authors:** Erika L Varner, Sophie Trefely, David Bartee, Eliana von Krusenstiern, Luke Izzo, Carmen Bekeova, Roddy S O’Connor, Erin L. Seifert, Kathryn Wellen, Jordan L. Meier, Nathaniel W. Snyder

## Abstract

Lysine lactoylation is a recently described protein post-translational modification (PTM). However, the biochemical pathways responsible for this acylation remain unclear. Two metabolite-dependent mechanisms have been proposed: enzymatic histone lysine lactoylation derived from lactoyl-coenzyme A (lactoyl-CoA, also termed lactyl-CoA), and non-enzymatic lysine lactoylation resulting from acyl-transfer via lactoyl-glutathione. While the former has precedent in the form of enzyme-catalyzed lysine acylation, the lactoyl-CoA metabolite has not been previously quantified in mammalian systems. Here we use liquid chromatography-high resolution mass spectrometry (LC-HRMS) together with a synthetic standard to detect and validate the presence of lactoyl-CoA in cell and tissue samples. Conducting a retrospective analysis of data from previously analyzed samples revealed the presence of lactoyl-CoA in diverse cell and tissue contexts. In addition, we describe a biosynthetic route to generate ^13^C_3_ ^15^N_1_ -isotopically-labeled lactoyl-CoA, providing a co-eluting internal standard for analysis of this metabolite. We estimate lactoyl-CoA concentrations of 1.14×10^−8^ pmol/cell in cell culture and 0.0172 pmol/mg tissue wet weight in mouse heart. These levels are similar to crotonyl-CoA, but between 20-350 times lower than predominant acyl-CoAs such as acetyl-, propionyl-, and succinyl-CoA. Overall our studies provide the first quantitative measurements of lactoyl-CoA and provide a methodological foundation for the interrogation of this novel metabolite in biology and disease.

**Highlights:**

- Detection of lactoyl-CoA at picomole concentrations across tissues and cells
- Lactoyl-CoA was detected at concentrations similar to crontonyl-CoA within HepG2 cells
- Isotopically labeled ^13^C_3_^15^N_1_-lactoyl-CoA can be prepared by SILEC

## Introduction

Acyl-coenzyme A (acyl-CoA) thioesters serve as critical metabolites in cellular energy metabolism. They also play a role in cellular signaling, acting as the acyl-donors for enzymatic and non-enzymatic PTMs collectively known as lysine acylation. Several studies have found that lysine acylation is sensitive to cellular metabolism, implying that acyl-CoA acyl-CoA levels may directly link metabolism to PTM-mediated signaling(1). This has the potential to occur either via the utilization of acyl-CoAs as rate-limiting cofactors in enzymatic protein modifications catalyzed by lysine acyltransferase (KAT) and histone acyltransfease (HAT) enzymes, or through non-enzymatic acylation reactions that can be heavily influenced by the electrophilicity of individual acyl-CoA species(2). A physiochemically diverse set of acylations have been detected on histone lysine residues, including not only the predominant acetylation but also succinylation, propionylation, (iso)butyrylation, crotonylation, malonylation, and several others(3). Lysine acylations generally correlate closely with the abundance of corresponding acyl-CoA(4).

Recently, Zhang et al reported a new addition to this emerging class of PTMs, known as lysine lactoylation (5). Applying antibody-based enrichment, this PTM was mapped to histone in M1 polarized macrophages and shown to correlate with altered gene expression during immune activation. Furthermore, reconstitution of lysine lactoylation within vitro transcription assays was found to be dependent on both the presence of a KAT enzyme (p300) as well as lactoyl-CoA, consistent with enzymatic acyl-transfer of the lactate group from the acyl-CoA to histones. Since lactate concentrations can shift from <1 mM to > 20mM across both normal and pathological processes (6), a biochemical link between lactate and histone modifications would be of interest in fields ranging from exercise physiology to cancer metabolism.

While microbial lactoyl-CoA production has been engineered in bio-industrial applications (7), the existence or quantification of lactoyl-CoA (also called lactyl-CoA and 2-hydroxypropanoyl-CoA) in mammalian cells or tissues has not been previously reported in the literature. Furthermore, the recent discovery of a parallel route to lysine lactoylaton via non-enzymatic acyl-transfer from lactoyl-glutathione raises the question as to whether these two pathways play distinct or similar roles (3). Since there is no described lactoyl-CoA synthetase or transferase that activates lactate to lactoyl-CoA in mammalian biochemistry, the in vitro and in vivo role of lactoyl-CoA remains poorly described. In order to better understand the potential role of enzyme-catalyzed lysine lactoylation in metazoans, here we report a method for the detection of lactoyl-CoA by liquid chromatography-high resolution mass spectrometry (LC-HRMS). Fortuitously, we find that lactoyl-CoA derivatives containing ^13^C_3_^15^N_1_-pantothenate are formed in human cells during stable isotope labeling by essential nutrients in cell culture (SILEC), providing access to an isotopically-labeled internal standard for quantitative studies (8). Applying this approach enables a comparative analysis of lactoyl-CoA and other metabolic acyl-CoAs, providing some initial insights into the concentration of this metabolite. By enabling the detection and quantitative analysis of lactoyl-CoA, our study provides the fundamental underpinnings for understanding the function of this metabolite in biology and disease.

## Methods

### Chemicals and Reagents

5-sulfosalicylic acid (SSA), trichloroacetic acid, and ammonium acetate were purchased from Sigma-Aldrich (St. Louis, MO). Optima LC-MS grade methanol (MeOH), acetonitrile (ACN), formic acid, and water were purchased from Fisher Scientific (Pittsburgh, PA).

### Synthesis of l-lactoyl-CoA

Synthesis was conducted as previously described with minor modifications (7). Briefly, l-lactic acid (90 mg, 1.0 mmol) and *N*-hydroxy succimide (NHS) were dissolved in anhydrous tetrahydrofuran (THF, 8 mL). A solution of dicyclohexylcarbodiimide (206 mg, 1.0 mmol) in THF (2 mL) was added dropwise to the l-lactic acid solution at 22 °C with stirring. The reaction mixture was stirred overnight at 22 °C. The resulting precipitate was removed by vacuum filtration and the filtrate was condensed under reduced pressure to yield l-lactyl-NHS which was used without further purification.

A solution of l-lactyl-NHS (9.3 mg, 50 mmol) in methanol (0.5 mL) was added to a solution of coenzyme A (25 mg, 33 mmol) in aqueous bicarbonate solution (0.5 mL, 500 mM, pH 8) and stirred at 22 °C for 3 h. The remaining aqueous solution was neutralized with trifluoroacetic acid (TFA, 20 mL) and immediately purified by C_18_-reversed phase-silica chromatography. Method: (solvents - A: 0.05% TFA in water, B: acetonitrile) gradient - 0-2 column volumes (CV), 0% B; 2-12 CV, 0-50% B; 12-13 CV, 50-100% B; and 13-25 CV, 100% B. Fractions containing the desired product were pooled and lyophilized to provide a fluffy white powder (7.3 mg, 26 % yield).

### Liquid chromatography-high resolution mass spectrometry

Acyl-CoAs were quantified by liquid chromatography-high resolution mass spectrometry as previously published (9, 10). For quantification from cells, media was aspirated from attached cells, 1 mL of ice cold 10% (w/v) trichloroacetic acid (Sigma Aldrich) in water was added, and cells were scraped and transferred to 1.5 mL Eppendorf tubes. Samples were spiked with 50 µL internal standard prepared as previously published (10) sonicated for 12 × 0.5 s pulses in, then protein was pelleted by centrifugation at 17,000 ×g from 10 min at 4 °C. The cleared supernatant was purified by solid-phase extraction using Oasis HLB 1cc (30 mg) SPE columns (Waters). Columns were washed with 1 mL methanol, equilibrated with 1 mL water, loaded with sample, desalted with 1 mL water, and eluted with 1 mL methanol containing 25 mM ammonium acetate. Samples were also extracted by addition of 1 mL −80°C 80:20 methanol:water, followed by sonication and centrifugation as above then evaporation of the supernatant. Retrospective analysis was also conducted on tissue samples using the extraction method of Minkler, *et al*. (11) modified for isotope dilution LC-MS (10). Calibration curves were prepared using analytical standards from Sigma Aldrich (for all acyl-CoAs other than lactoryl-CoA) and processed identically to the samples.

The purified extracts were evaporated to dryness under nitrogen then suspended in 55 μl 5% (w/v) 5-sulfosalicylic acid in optima HPLC grade water. 5 μl of samples in 5% SSA were analyzed by injection of an Ultimate 3000 HPLC coupled to a Q Exactive Plus (Thermo Scientific) mass spectrometer in positive ESI mode using the settings described previously (9), or for the full acyl-length extraction (12). Modifications were made to these methods for MS/HRMS experiments as described in each experiment for acquisition of precursor and product ions. Data were integrated using XCalibur Quan and Qual Browsers (Thermo Scientific), Tracefinder v4.1 (Thermo Scientific) software, and additional statistical analysis conducted by Prism v7.05 (GraphPad).

### Cell culture

Cells were passaged every 2–3 days at 80% confluence. Human hepatocellular carcinoma (HepG2) cells were used at <20 passages from ATCC stocks (ATCC) and were cultured in high glucose DMEM (Thermo-Fisher Scientific), supplemented with 10% fetal bovine serum (Gemini Biosciences) and penicillin/streptomycin (Thermo Fisher Scientific). All cells were tested mycoplasma-free.

### Heart and muscle tissue analysis

Experiments were performed on 11–13-week-old male and female C57BL/6J mice. Mice were maintained on a 12-12 h light-dark cycle (lights on: 7:00 AM to 7:00 PM) and ad libitum fed a standard diet (LabDiet 5001, Purina). Fasted mice were fasted overnight (7:00 PM to 9:00 AM). Whole heart samples were always harvested in the morning, snap frozen, and stored at −80°C. All protocols were approved by the Thomas Jefferson University Institutional Animal Care and Use Committee (protocol number IACUC 01307).

## Results

### Synthesis of lactoyl-CoA

To define detection and identification of lactoyl-CoA we synthesized a pure standard. The resulting product was a white crystalline solid. Both NMR and LC-HRMS of the pure standard provided evidence of the structure of lactoyl-CoA (**Fig. 1**). ^1^H NMR (500 MHz, D_2_O) δ 8.56 (d, *J* = 5.7 Hz, 1H), 8.35 (s, 1H), 6.13 (d, *J* = 5.2 Hz, 1H), 4.81 (s, 2H), 4.52 (s, 1H), 4.29 (q, *J* = 7.0 Hz, 1H), 4.19 (s, 2H), 3.94 (s, 1H), 3.79 (d, *J* = 9.3 Hz, 1H), 3.54 (d, *J* = 9.2 Hz, 1H), 3.37 (t, *J* = 6.7 Hz, 2H), 3.26 (t, *J* = 6.5 Hz, 2H), 2.35 (t, *J* = 6.7 Hz, 2H), 1.26 (d, *J* = 7.0 Hz, 3H), 0.86 (s, 3H), 0.74 (s, 3H). For HRMS, we observed an intense [MH]+ ion (m/z = 840.1435) matching the theoretical m/z calculated for C_24_H_41_N_7_O_18_P_3_S+ (840.1436138) (Δppm= −0.135). Based on previous literature of the stability of acyl-CoAs in acid and lowered temperature avoiding excessive freeze thaw cycles (10) we re-suspended the product in 10% TCA in water at 1 mg/ mL to generate aliquots of concentrated stocks for future use.

**Figure 1.**
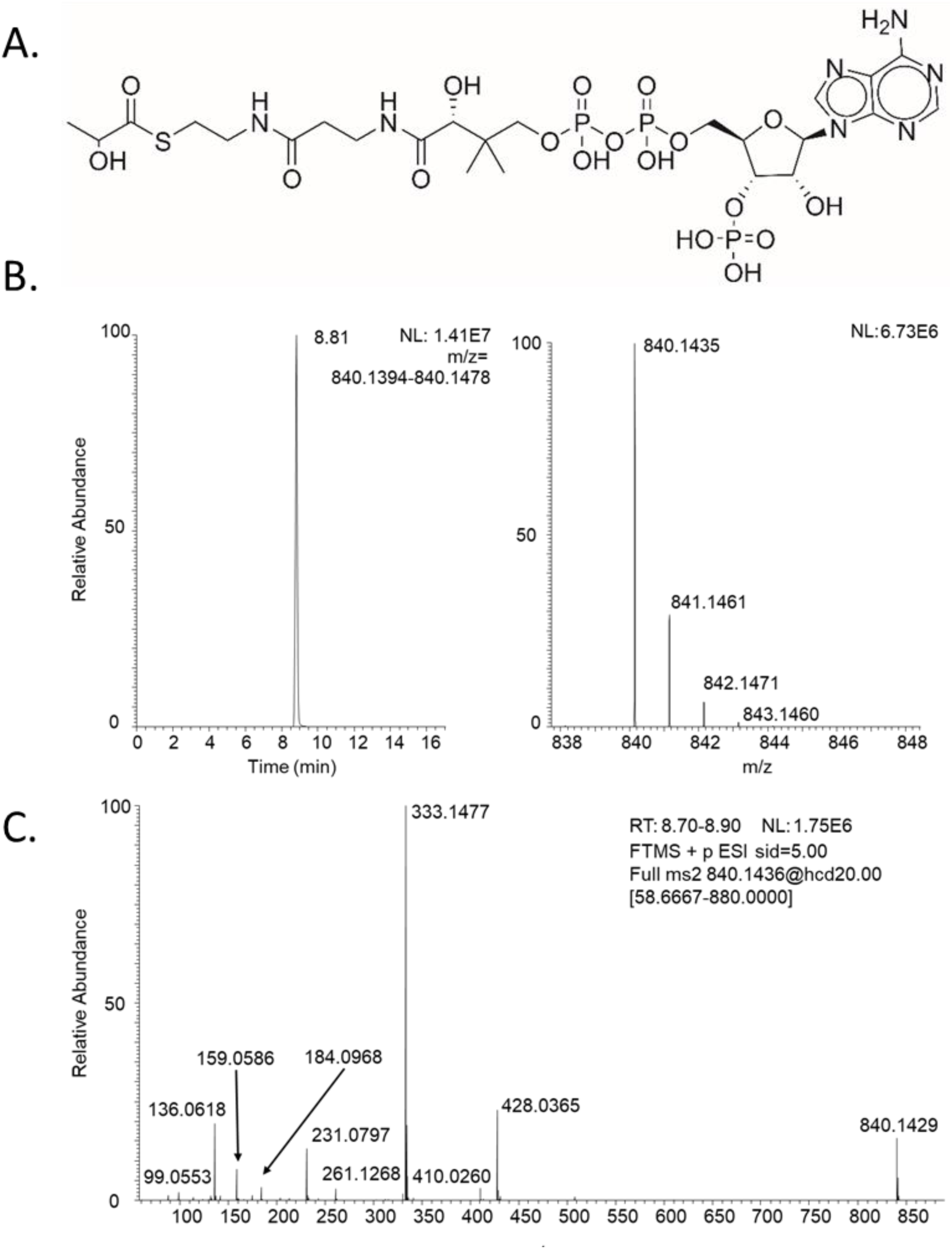
(A) Structure of lactoyl-CoA (stereochemistry not shown), (B) LC-HRMS of synthetic lactoyl-CoA, (C) LC-MS/HRMS of lactoyl-CoA.

### Qualitative characterization of lactoyl-CoA in cells and tissues

Using LC-HRMS and LC-MS/HRMS we detected lactoyl-CoA from cells. The cell derived lactoyl-CoA matched the synthetic standard on retention time, HRMS and MS/HRMS on major product ions (**Fig. 2**). In contrast, extraction of acyl-CoAs from a different 10 cm plate of cells with protein precipitation by −80°C 80:20 methanol: water did not yield a discernable peak above noise (data not shown). The retention time of the putative lactoyl-CoA (tr= 8.8 min) in reversed phase chromatography fell earlier than propionyl-CoA (tr = 9.2) corresponding to the additional hydroxyl group in the acyl-group.

**Figure 2.**
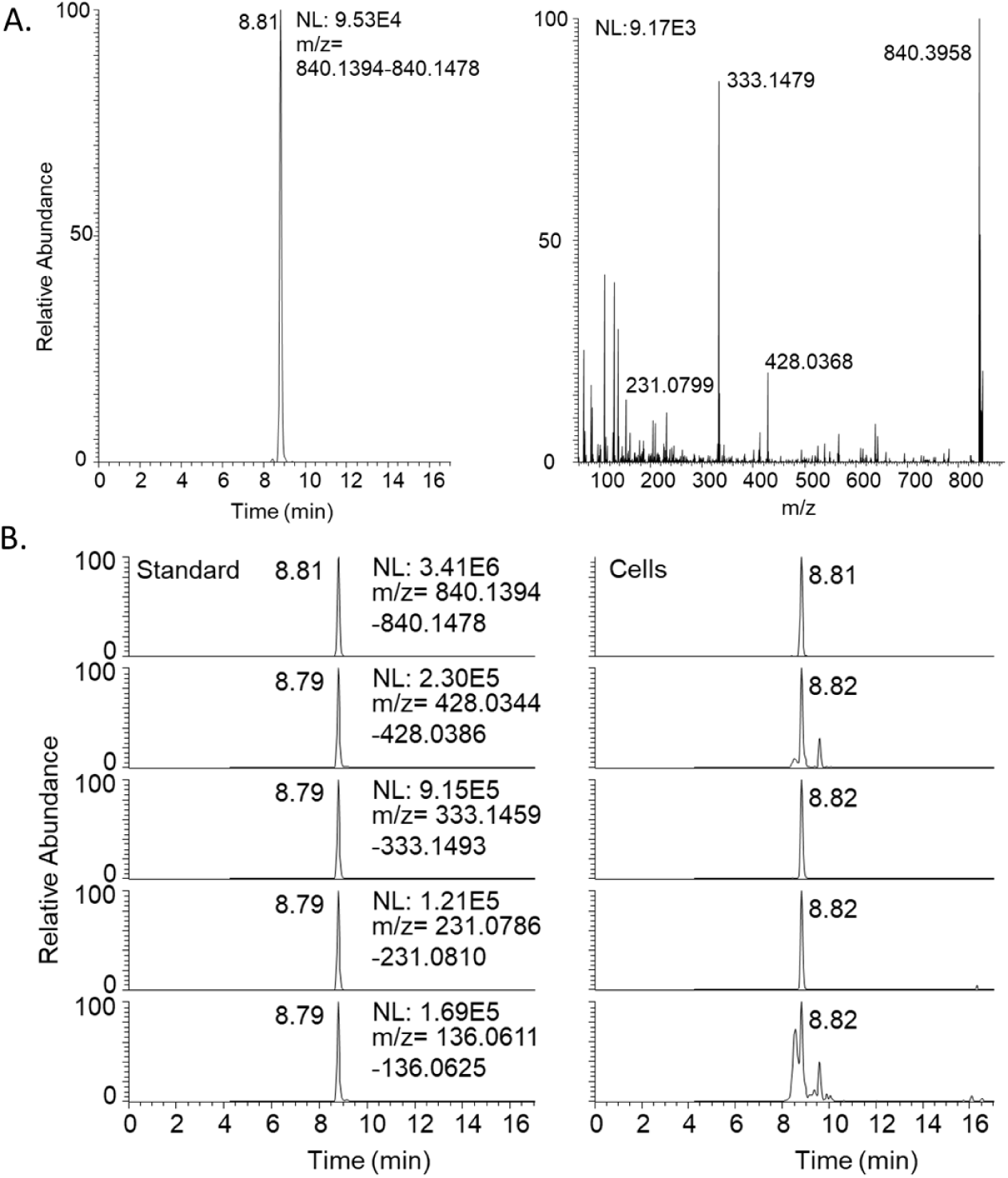
(A) LC-HRMS of lactoyl-CoA from HepG2 cell extract and (B) LC-MS/HRMS of synthetic lactoyl-CoA (left) and the same ions in cell extract (right).

Since acyl-CoAs produce a wide variety of product ions upon collision induced dissociation, we examined the likely specificity of each major product ion via plotting chromatograms for the top 4 ions derived from lactoyl-CoA (**Fig. 2**). Within cells, the product ion corresponding to the M-507 (M-506.99575) neutral loss (m/z 333.14786) was the most intense and specific.

Based on the reanalysis of data from samples previously acquired by LC-HRMS, we examined the detection of lactoyl-CoA in other cellular and tissue contexts. Since our routine methods of acyl-CoA analysis include a full scan acquisition covering the [MH]+ ion we predominantly detected from the synthetic standard, this allowed semi-quantitative retrospective analysis of previously acquired data. Additional experiments using a mixed chain length acyl-CoA extraction from Minkler, *et al*. (11) similarly detected a LC-HRMS and LC-MS/HRMS peak corresponding to lactoyl-CoA from heart and muscle samples previously acquired, which we then matched to a standard acquired with the same chromatographic method (**Fig. 3**).

**Figure 3.**
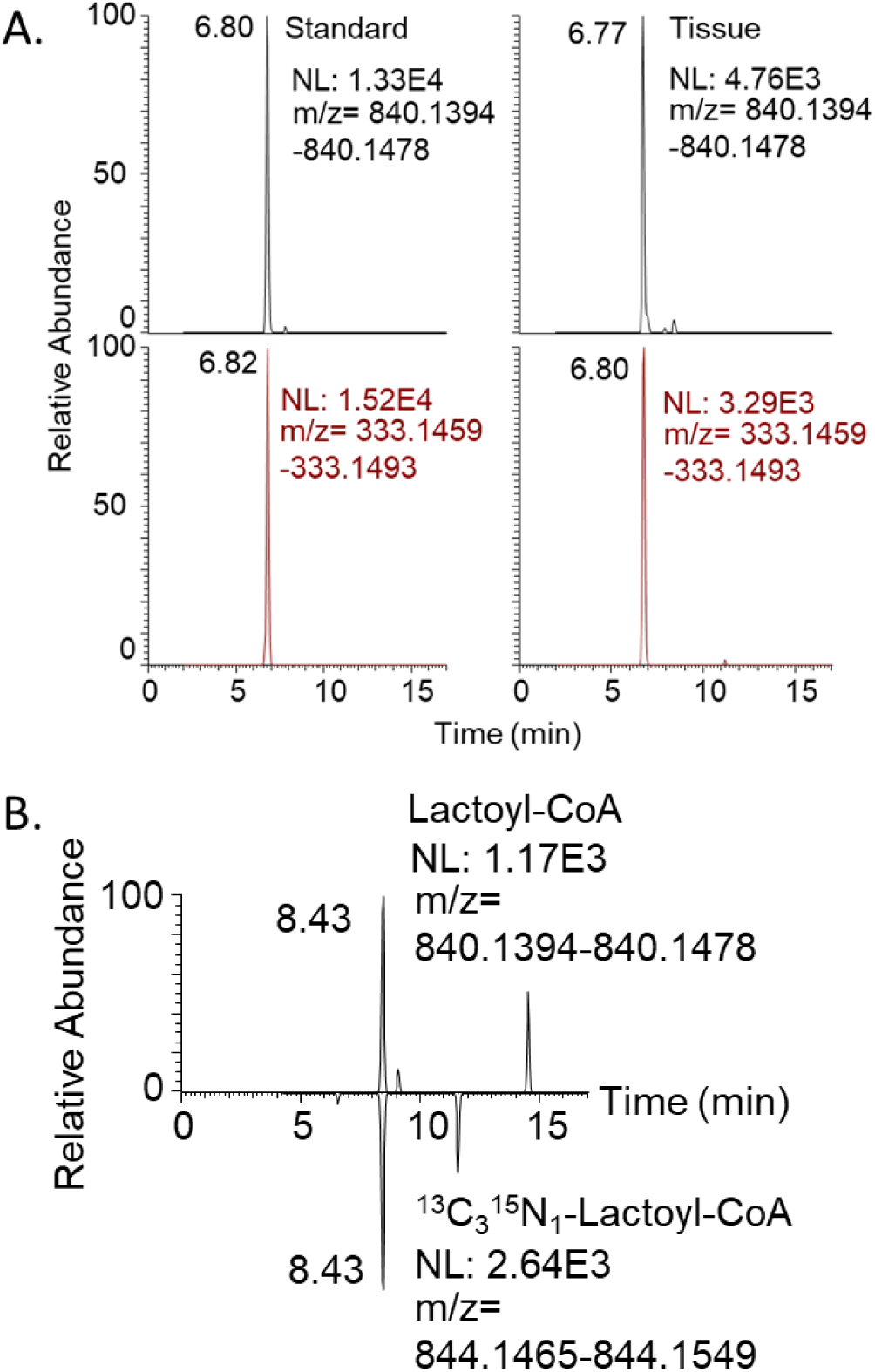
Detection of lactoyl-CoA from retrospective analysis (A) of tissues by LC-HRMS (top) and LC-MS/HRMS (bottom) and (B) co-elution of lactoyl-CoA with ^13^C_3_^15^N_1_-lactoyl-CoA from retrospective data.

From previous stable isotope labeling using essential nutrients in cell culture where acyl-CoA libraries incorporating ^13^C_3_ ^15^N_1_ -pantothenate at high efficiency (>99.5%) were generated and spiked into samples, we interrogated generation and co-elution of ^13^C_3_ ^15^N_1_ -labeled lactoyl-CoA with unlabeled lactoyl-CoA (**Fig. 3**). Previous studies using pantothenate SILEC in HepG2 cells demonstrated co-elution of lactoyl-Coa with ^13^C_3_ ^15^N_1_ - labeled lactoyl-CoA. However, we found unfortunately that the production of acyl-CoA internal standards by yeast SILEC in the conditions we commonly used did not produce detectable amounts of ^13^C_3_ ^15^N_1_ -labeled lactoyl-CoA. This lack of ^13^C_3_ ^15^N_1_ -lactoyl-CoA in these samples strongly suggests that there is no post-extraction acyl-transfer of lactate to CoASH, as the SILEC standard includes ^13^C_3_ ^15^N_1_ -CoASH that would have artificially formed ^13^C_3_ ^15^N_1_ -lactoyl-CoA if possible.

### Partial re-validation of a quantitative method to include lactoyl-CoA

Extending previously published quantitative methodology for acyl-CoA quantification for short- and longer chain acyl-CoAs(9), we partially revalidated the LC-HRMS method for select quantitative properties of lactoyl-CoA. Linear range was estimated for isotope dilution and label free quantification. In both cases, lactoyl-CoA was linear (R^2^ > 0.998) and observed values within 25% of expected values from 0.03 pmol per sample (3 fmol on column) to the highest range tested at 500 pmol per sample (50 fmol on column). Lactoyl-CoA was detectable down to 0.002 pmol per sample (0.2 fmol on column), but quantification was unstable across runs over 24 hours. Re-injection of samples from both purified standards, and purified standards spiked into cellular extract were stable (+/- 5%) over 24 hours in a chilled autosampler. Unfortunately, as noted, our method of producing internal standards in yeast did not generate a lactoyl-CoA under the conditions we used. Thus, we investigated using surrogate internal standards including ^13^C_3_^15^N_1_-acetyl-, ^13^C_3_ ^15^N_1_-succinyl- and ^13^C_3_ ^15^N_1_-HMG-CoA. All produced linear calibration curves with high R^2^-values (> 0.998) when plotting the ratios of analyte/each internal standard as the dependent variable and similar deviation of calibration points from observed versus expected values.

### Lactoyl-CoA is in low abundance relative to other commonly quantified short chain acyl-CoAs

We quantified lactoyl-CoA via relative response factor to other short chain acyl-CoAs for retrospective sample analysis. This allowed us to rank the molar amount of lactoyl-CoA versus other acyl-CoAs detected in cells. We calculated proportionality constants using known amounts of lactoyl-CoA across a calibration curve of serial dilutions of cell and tissue relevant concentrations of other short chain acyl-CoAs. Proportionality constants on a molar basis were calculated as:

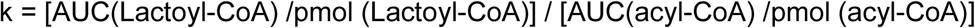

Propionyl-CoA and succinyl-CoA had the response factors that most closely approximated 1, where the same molar amount of lactoyl-CoA and other acyl-CoAs generated the same intensity of signal. Thus, we used propionyl-CoA as a surrogate to retrospectively estimate lactoyl-CoA concentrations in a semi-quantitative manner. We found that lacotyl-CoA in HepG2 cells was quantifiable at 0.011 pmol/10^6 cells, comparable to crontonyl-CoA (0.033 pmol/10^6 cells), but markedly less abundant (30-250 times lower) than major acyl-CoAs such as acetyl-, succinyl-, and propionyl-CoA (**Table 1**).

**Table 1.** Concentrations of short chain acyl-CoAs within HepG2 cells.

| Acyl-CoA | pmol/10<sup>6</sup> cells | Standard Deviation |
| --- | --- | --- |
| Acetyl-CoA | 10.644 | 1.364 |
| Succinyl-CoA | 25.467 | 2.818 |
| Propionyl-CoA | 3.532 | 0.652 |
| CoASH | 1.734 | 0.189 |
| Butyryl-CoA | 1.013 | 0.159 |
| HMG-CoA | 0.971 | 0.326 |
| Glutaryl-CoA | 0.647 | 0.112 |
| Valeryl-CoA | 1.118 | 0.143 |
| Crotonoyl-CoA | 0.032 | 0.015 |
| Lactoyl-CoA | 0.011 | 0.003 |

Since we could also detect lactoyl-CoA in tissues, we performed the same semi-quantitative estimation in heart tissue from mice. In these experiments, since we did not have ^13^C_3_^15^N_1_-lactoyl-CoA due to the method of production of the internal standard, we also used ^13^C_3_^15^N_1_-propionyl-CoA as a surrogate internal standard based on the similarity in response generated from equimolar amounts found above. We compared the lactoyl-CoA concentration of murine heart by sex and by fed/fasted status. Across 25 mouse samples of murine heart we retrospectively estimated a mean (standard deviation) of 0.0179 (0.0153) pmol/mg tissue wet weight. We could detect no difference between fasted (0.0187 pmol/mg) or fed (0.0172 pmol/mg). Similarly, we found no difference between lactoyl-CoA content in hearts from male [0.0214 (0.0173) pmol/mg] and female [0.0145 (0.0119)) mice in this experiment. In these same tissues, acetyl-CoA was 5.77 (3.08) pmol/mg and propionyl-CoA 0.476 (0.224) pmol/mg. Thus, again we found lactoyl-CoA to be 335 (vs acetyl-) and 27 (vs propionyl-) times lower than major short chain acyl-CoAs, but still detectable.

## Discussion

Increasingly sensitive proteomics approaches discover and describe a large abundance of protein PTMs. These PTMs are commonly putatively attributed to acyl-transfer from various metabolites including biologically activated or reactive metabolite classes including acyl-CoA thioesters (1). Due to the shared physiochemical properties of these classes of metabolites, methods of purification, separation, and analysis for certain members of these classes of metabolites tend to also capture other members of the same metabolite family. This becomes increasingly evident with methods of detection including HRMS that can use less restrictive analyte detection leading to a rise of a combination of targeted and untargeted metabolomics, termed hybrid metabolomics where a limited targeted set of analytes is profiled, but metabolomics level interrogation on the background data can still be performed (13). As we demonstrate here, the re-interrogation of previously acquired background metabolomics data can be useful for metabolite discovery and semi-quantification. This was especially useful because our approach to internal standardization using the SILEC approach, generated a library of stable isotope labeled internal standards that fortuitously included ^13^C_3_^15^N_1_-lactoyl-CoA. Other approaches to internal standard production from isotope labeled biological sources may also show this same benefit (14). However, the yeast-based method of SILEC production used here did not produce detectable ^13^C_3_^15^N_1_-lactoyl-CoA which is unfortunate for quantification purposes but does indicate that acyl-transfer post-extraction did not result in artifactual lactoyl-CoA generation. Such an artifact is certainly not unprecedented, as acyl-groups are known to migrate to and from CoASH, and transacylation of carboxyclic acid containing acyls-groups is biologically and pharmacologically common (15).

Our data does suggest that lactoyl-CoA generation is relatively less abundant than other acyl-CoAs. This is unsurprising as histone lysine lactoylation was reported in conditions including hypoxia and M1 macrophage polarization where lactate production is high(5). Thus, there is likely a precedent for contexts where lacotyl-CoA would be more abundant. The quantitative comparison to crontonyl-CoA, which is also thought to be the acyl-donor for histone crontoylation, may put these modifications in the same realm of biological plausibility(16). However, our retrospective analysis did not detect high levels of lactoyl-CoA in conditions where we expected it. This included cell culture within incubation of exogenous lactate in T-cell culture (data not shown). The reasons for this discrepancy across detection require systematic investigation in the future. Similarly, investigations of the comprehensive changes of other acyl-CoAs (including CoASH) that could compete with lactoyl-CoA for formation of acyl-transfer to lysine would be warranted.

This study has two important limitations that warrant discussion. First, although the description of lyisine lactoylation was the reason we synthesized then searched for lactoyl-CoA this study does not quantify either lactoyl-lysine or lacotyl-GSH. Future studies will be needed to compare the relative concentrations of each of these acylated molecules and the biological contexts where they may be important. As noted recently by Kulkarni and Brookes, Kla would be isobaric to *N*^*6*^-carboxyethyl lysine resulting from methyglyoxal, a reactive side-product of glucose metabolism (17). Thus, future studies should resolve these potentially interfering species via isotope tracing or detection methods capable of discerning the two lysine modifications. Second, our methods were operating near their estimated limits of detection for lactoyl-CoA which poses problems of censoring and imposes limits on experimental design. Further improvements in analysis, including more sensitive LC-MS approaches reaching atto- and zepto-mole on column quantification may be needed considering the low abundance of lactoyl-CoA. Increased sensitivity would be especially necessary in isotropic tracing applications where additional sensitivity is required since labeling distributes signal intensity across isotopologues. Such isotope tracing experiments from labeled glucose and lactate would be one of the most straightforward ways of establishing the route and mechanism of lactoyl-lysine formation and are tractable even in human studies (18). Additionally, future methods could achieve separation of enantiomeric acyl-CoAs, including lactoyl-CoA (which could form from d- or l-lactic acid). These analytical improvements would be useful in the next steps of elucidating the as yet unknown biological function, enzymology, and occurrence of lactoyl-CoA.

## Conclusion

We can detect and quantify lactoyl-CoA within cells and tissues at low pmol abundance. This should be valuable for future studies interrogating the role of lactoyl-CoA and its mechanisms of production and use within wider biologic contexts. The utility of LC-HRMS data for retrospective reanalysis to examine newly described metabolites also highlights the power of this approach over more targeted acquisition strategies.

## Acknowledgements

This work was funded by R01GM132261 to N.W.S, the Intramural Research Program of NIH, the National Cancer Institute, The Center for Cancer Research (ZIA BC011488-06) to J.L.M., ADA fellowship 1-18-PDF-144 to S.T., T32GM-07229 to L.I., R01CA228339 to K.W., and R01DK109100 to E.L.S.

## Notes

### Competing Interest Statement

The authors have declared no competing interest.

### Summary of Updates

Additional discussion and results have been added.

